# Photosynthetic conversion of CO_2_ to hyaluronic acid by engineered cyanobacteria

**DOI:** 10.1101/691543

**Authors:** Lifang Zhang, Tiago Toscano Selão, Peter J. Nixon, Birgitta Norling

## Abstract

Hyaluronic acid (HA), consisting of alternating N-acetylglucosamine and glucuronic acid units, is a natural polymer with diverse cosmetic and medical applications. Currently, HA is produced by overexpressing HA synthases from gram-negative *Pasteurella multocida* (encoded by *pmHAS*) or gram-positive *Streptococcus equisimilis* (encoded by *seHasA*) in various heterotrophic microbial production platforms. Here we introduced these two different types of HA synthase into the fast-growing cyanobacterium *Synechococcus* sp. PCC 7002 (Syn7002) to explore the capacity for producing HA in a photosynthetic system. Our results show that both HA synthases enable Syn7002 to produce HA photoautotrophically, but that overexpression of the soluble HA synthase (PmHAS) is less deleterious to cell growth and results in higher production. Genetic disruption of the competing cellulose biosynthetic pathway increased the HA titer by over 5-fold (from 14 mg/L to 80 mg/L) and the relative proportion of HA with molecular mass greater than 2 MDa. Introduction of *glmS* and *glmU*, coding for enzymes involved in the biosynthesis of the precursor UDP-N-acetylglucosamine, in combination with partial glycogen depletion, allowed photosynthetic production of 112 mg/L of HA in 5 days, an 8-fold increase in comparison to the initial PmHAS expressing strain. Addition of *tuaD* and *gtaB* (coding for genes involved in UDP-glucuronic acid biosynthesis) also improved the HA yield, albeit to a lesser extent. Overall our results have shown that cyanobacteria hold promise for sustainable production of pharmaceutically important polysaccharides from sunlight and CO_2_.

## 1. Introduction

Cyanobacteria are gaining attention as hosts for photosynthetic production of high-value molecules due to their easier genetic manipulation in comparison to other photosynthetic systems and favourable growth rates [1, 2]. Metabolic engineering of cyanobacteria has successfully led to the production of a wide range of industrially relevant products [3, 4]. However, apart from a very recent report on the production of heparosan [5], the use of cyanobacteria to produce pharmaceutically important polysaccharides remains relatively unexplored.

Hyaluronic acid (HA) is a unique biopolymer composed of alternating β-1,3-N-acetyl glucosamine and β-1,4-glucuronic acid disaccharide units [6]. Its distinctive moisturizing and viscoelastic properties, coupled to a lack of immunogenicity and toxicity, have led to a wide range of proven and marketed applications for HA within the cosmetic and biomedical industries [7]. The global HA market was valued at USD 7.2 billion in 2016 and is expected to reach USD 15.4 billion by 2025 [8]. However, most of the commercial HA is currently either isolated from animal sources, such as rooster combs or made by microbial synthesis via attenuated strains of group C *Streptococci* [9], with the potential problem of contamination by endotoxins. The increased demand for HA and arising safety concerns have led researchers to use synthetic biology and metabolic engineering approaches to develop alternative sources of HA production.

Two types of HA synthase, catalyzing the final HA synthesis step, are commonly used in metabolic engineering efforts, most commonly the enzymes PmHAS from *Pasteurella multocida* and SeHasA (or SzHasA) from *Streptococcus* species. Successful HA production was previously demonstrated in genetically modified non-pathogenic *Bacillus subtilis* and *Escherichia coli* strains overexpressing different HA synthases, along with the relevant precursor synthesis enzymes [10-14]. Although HA titers in these modified strains are promising, with some being used in industrial scale processes [14], their production is completely based on fermentative processes, requiring the input of substantial amounts of carbon feedstocks, such as glucose or sucrose. On the other hand, the photosynthetic process in cyanobacteria converts CO_2_ into carbohydrates, and some strains have been engineered to efficiently produce and excrete either soluble sugars (such as sucrose [15]) or polysaccharides, e.g. crystalline cellulose [16].

In this study, the chassis strain *Synechococcus* sp. PCC 7002 (hereafter Syn7002) was engineered into a photosynthetic HA production system by introducing different HA synthases and improvements to HA production were further investigated by enhancing precursor biosynthesis pathways and/or blocking some of the competing pathways. Our results show that cyanobacteria can be manipulated to produce hyaluronic acid from CO_2_, demonstrating that they have the potential to become viable alternatives for the sustainable production of biomedically relevant polysaccharides.

## 2. Material and Methods

### 2.1. Strains and culture conditions

Wild-type *Synechococcus* sp. PCC 7002 (a kind gift from Donald Bryant, Pennsylvania State University, USA) and all derivative strains were grown in medium AD7 [17] and supplemented with antibiotics as required, namely chloramphenicol (10 μg/mL) and/or kanamycin (100 μg/mL). Liquid cultures and plates were grown in an atmosphere of CO_2_-enriched (1% (v/v)) air, under continuous illumination (300 μmol photons m^−2^ s^−1^) at 38 °C, as previously described [17]. Growth was monitored by measuring OD_730_ in a plate reader (Hidex Sense, Hidex) and utilizing an in-house generated 70-point calibration curve (R^2^=0.9818) converting OD_730_ in the plate reader to that measured using a 1 cm light path table top spectrophotometer (Cary 300Bio, Varian), using AD7 as blank. Dry cell weight at 5 days was measured as previously described [17]. All growth measurements were performed using biological triplicates (n=3). For HA production test, cultures were pre-adjusted to an OD_730_=1, induced by addition of 1 mM IPTG, and samples collected at the time points indicated. Cell pellets and supernatants were stored at −20 °C until further use.

### 2.2. Cyanobacterial strain construction

Genes encoding two HA synthases, *pmHAS* and *seHasA* (with a C-terminal FLAG tag), as well as *B. subtilis* UDP-glucose dehydrogenase (*tuaD*), UDP-glucose pyrophosphorylase (*gtaB*), and the *E. coli* enzymes glutamine amidotransferase (*glmS*) and acetyl-CoA acetyltransferase/pyrophosphorylase (*glmU*, a homologue of *gcaD* in *B. subtilis*) were codon-optimized for Syn7002 and synthesized by GenScript (Hong Kong). The expression of both HA synthase genes was controlled by an IPTG inducible promotor, P_cLac143_, based on plasmid *pAcsA-*P_cLac143-_*YFP* (a kind gift from Brian Pfleger, University of Wisconsin-Madison, USA) [18]. The *tuaD* and *gtaB* genes were synthesized as an artificial operon, under control of the strong constitutive P_cpc560_ promoter [19], with a second artificial operon containing *glmS* and *glmU*, controlled by the strong constitutive P_cpt_ promoter [20]. In all cases, the strong AGGAGA RBS sequence was utilized, with a random 8 bp DNA sequence between RBS and the initial ATG codon [20]. All enzymes utilized were purchased from NEB unless otherwise specified and all primers used are listed in Table S1. DNA fragments were PCR amplified with Q5 DNA polymerase, purified using the EZ-10 Spin Column PCR Products Purification Kit (Bio-Basic, Canada) and assembled either using the pEASY-Uni Seamless Cloning and Assembly Kit (TransGen Biotech, China), following manufacturer’s instructions, or by *E. coli* mediated assembly [21]. Supercompetent *E. coli* cells (Stellar, TaKaRa) were used for all cloning steps and were grown in LB medium supplemented with antibiotics as required - ampicillin (100 μg/mL), chloramphenicol (25 μg/mL) or kanamycin (50 μg/mL). The sequence of all constructed plasmids (Table S1) was confirmed by Sanger sequencing. Syn7002 transformation was performed according to standard protocols [22]. Full genomic segregation in modified strains was confirmed by colony (gDNA) PCR using specific primers. A list of all strains generated in this work is presented in Table 1.

**Table 1.** Engineered cyanobacterial strains used in the study.

| Strain name | Genotype | Characteristics |
| --- | --- | --- |

**Table 1.** Engineered cyanobacterial strains used in the study.
|  |  |  |
| --- | --- | --- |
| HA01(HA02) | $\Delta AcsA::P_{cLac143} -pmHAS(-sfGFP)$ | HA producing, without or with sfGFP |
| HA03(HA04) | $\Delta AcsA::P_{cLac143} -seHasA(-sfGFP)$ | HA producing, without or with sfGFP |
| HA06 | $\Delta glgA2::Km^R + \Delta AcsA::P_{cLac143} -pmHAS$ | Partial glycogen depletion, HA producing |
| HA07 | $\Delta glgA1::Cm^R + \Delta AcsA::P_{cLac143} -pmHAS$ | Partial glycogen depletion, HA producing |
| HA08 | $\Delta cesA::Cm^R + \Delta AcsA::P_{cLac143} -pmHAS$ | Cellulose depletion, HA producing |
| HA11 | $\Delta glpK::P_{cpt} -glmS-glmU-Cm^R + \Delta AcsA::P_{cLac143} -pmHAS$ | Overexpression of heterologous UDP-GlcNAc pathway enzymes, HA producing |
| HA12 | $\Delta glgA1::P_{cpt} -glmS-glmU-Cm^R + \Delta AcsA::P_{cLac143} -pmHAS$ | Overexpression of heterologous UDP-GlcNAc pathway enzymes, partial glycogen depletion, HA producing |
| HA13 | $\Delta glgA2::P_{cpc560} -tuaD-gtaB-Km^R + \Delta AcsA::P_{cLac143} -pmHAS$ | Overexpression of heterologous UDP-GlcUA pathway enzymes, partial glycogen depletion, HA producing |

### 2.3. Fluorescence microscopy and western blot

Preparation of Syn7002 cells for confocal microscopy was performed as described earlier [23]. Essentially, cells from IPTG-induced cultures were collected and blotted on a 1% agarose gel pad (prepared with AD7 medium) for image analysis. Laser scanning confocal microscopy was performed using an LSM710 (Carl Zeiss), with a 1.4 NA Plan-Apo 60x oil immersion lens used as an objective at a zoom factor of 10 and excitation at 488 nm. Image analysis was carried out using the Zen software (Carl Zeiss, version 2.3). Cell size estimates were made using the measuring tool in the Zen software and calculating averages and standard deviations from 15-20 cells from each strain, imaged at the same magnification.

Whole-cell lysates were isolated as previously described and subjected to western blot analysis [24]. 10 µg total proteins were separated on 12% precast SDS-PAGE gels (BioRad), transferred onto PVDF membranes and probed with primary antibodies raised in rabbit against a synthetic PmHAS peptide (NDNDLKSMNVKGAS, amino acids 860 to 874, prepared by GenScript, Hong Kong) and/or a monoclonal mouse anti-FLAG M2 antibody (F3165, Sigma-Aldrich).

### 2.4. Cyanobacterial HA preparation and quantification by ELISA

HA production by engineered Syn7002 strains was routinely assessed by measuring the HA released into the growth medium at the indicated time points. Cells were collected by centrifugation (3000 g, 10 min, room temperature) and the cleared supernatant was used for HA assay. To concentrate HA from cyanobacterial culture, 3 volumes of ice-cold ethanol were added to the cleared supernatants and the resulting pellet was dried in air and re-suspended in deionised H_2_O for further use [25]. To evaluate the HA secretion ability of different strains, released HA (R-HA), capsular HA (bound to the cell surface, CPS-HA) and intracellular HA (intra-HA) were isolated according to previously described methods [10, 26], with whole cell lysates for intra-HA quantification prepared according to Selão et al [24]. HA quantification was performed using a Hyaluronan Quantikine ELISA Kit (DHYAL0, R&D systems, USA), following manufacturer’s instructions, either directly or by diluting samples with AD7 prior to quantification, and absorbance of the ELISA strips was measured using the Hidex Sense plate reader.

### 2.5. Characterization of HA by HPLC, LC-MS/MS and FTIR

Cyanobacterial HA was further analysed using an UFLC Prominence (Shimadzu, Japan) equipped with a size-exclusion chromatography column (Shodex OHpak, SB806 M HQ, 8.0 mm×300 mm, 13µm particle size, Shimadzu), coupled to a UV detector (Shimadzu, Japan) and a differential refractive index detector (dRI, Optilab rEX, Wyatt Technology, USA). The mobile phase was a 0.1 M NaCl solution and all chromatography runs were performed at 0.5 mL/min. Samples were filtered through 0.2 µm Whatman Puradisc 13 syringe filters (GE Healthcare, USA) prior to injection and 150 µL were used in each run. Chromatography data was recorded and analyzed by the ASTRA software (version 5.3, Wyatt Technology, USA). All peaks detected by the dRI detector were collected and saved at −20 °C until further use. The molecular mass of detected peaks was estimated by comparison to the elution times of PEG analytical standard (ReadyCal Set, #02393, Sigma-Aldrich) and of commercial HA with known molecular mass (2-2.2 MDa, #51967, Sigma-Aldrich).

LC-MS/MS was used to characterize the forming units of HA polymer produced by the PmHAS-expressing strains following hyaluronidase digestion. HPLC-purified sample was precipitated using ethanol as described above, and subjected to hyaluronidase (#H3506, Sigma-Aldrich) digestion according to manufacturer’s instructions. After confirmation of complete digestion using both the HA ELISA assay and HPLC, the resulting fragments were analyzed by LC-MS/MS, using a Xevo-TQ-S (Waters, Milford, USA) mass spectrometry system, coupled to an ACQUITY UPLC system (Waters) with a 2.1mm × 100mm HSS T3 Column, 1.8µm particle size (Waters), as previously described [27]. Cyanobacterial HA from HPLC eluted peaks was also subjected to Fourier transform infrared spectroscopy (FTIR) analysis after precipitation with cold ethanol. 10 µL samples were evenly spotted on a horizontal ZnSe ATR element (Pike Technologies) and dried under a stream of dry, CO_2_-free air. The respective spectra (average from 200 scans at 4 cm^−1^ resolution) were recorded between 4000-650 cm^−1^ using an FTIR Nicolet Nexus 470 instrument (Thermo Scientific), purged with dry, CO_2_-free air and equipped with an MCT/A detector cooled with liquid nitrogen.

## 3. Results and Discussion

### 3.1. Overexpression of HA synthase in Syn7002

As with all cyanobacteria sequenced so far, Syn7002 lacks homologues of the known HA synthases and is not known to naturally produce HA. However, an analysis of its predicted metabolic network (using the KEGG database) suggests that this organism has all the enzymes required to synthesize the two HA precursor molecules, UDP-GlcUA and UDP-GlcNAc (Fig. 1). Both of these UDP-sugars, as well as their precursors, are used for cyanobacterial cell wall biosynthesis and could theoretically be redirected to the biosynthesis of HA or similar polysaccharides [5].

**Figure 1.**
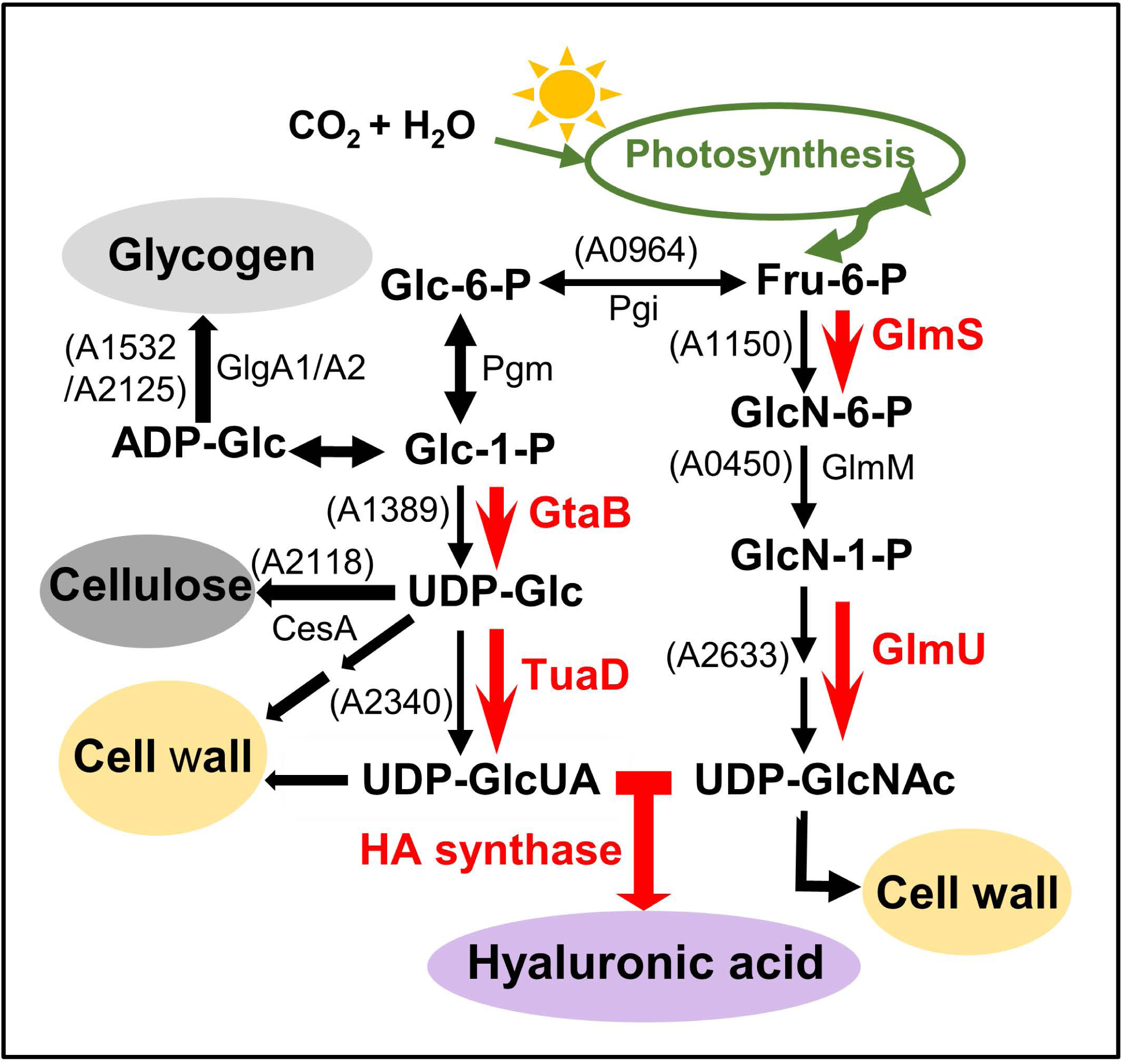
Schematic pathway for hyaluronic acid precursor biosynthesis in Syn7002 (adapted from [14]. Red arrows indicate heterologous enzymes commonly used to strengthen the precursors biosynthesis and HA formation in heterotrophic bacteria; equivalent, functionally proven or predicted enzymes from Syn7002 are shown in brackets. Selected potential competing pathway, such as glycogen, cellulose or cell wall biosynthesis are also marked with coloured-in circles. Glc-6-P: glucose-6-phosphate; Glc-1-P: glucose-1-phosphate; ADP-Glc: ADP-glucose; UDP-Glc: UDP-glucose; UDP-GlcUA: UDP-glucuronic acid; Fru-6-P: fructose-6-phosphate; GlcN-6-P: glucosamine-6-phosphate; GlcN-1-P: glucosamine-1-phosphate; GlcNAc-1-P: N-acetylglucosamine-1-phosphate; UDP-GlcNAc: UDP-N-acetylglucosamine; TuaD: UDP-glucose dehydrogenase; GtaB: UDP-glucose pyrophosphorylase; GlmS: glutamine amidotransferase; GlmU: acetyl-CoA acetyltransferase/pyrophosphorylase; GlgA1/A2: glycogen synthase A1/A2; CesA: cellulose synthase; Pgi: phosphoglucoseisomerase; Pgm: phosphoglucomutase

We used a previously described IPTG-inducible expression system (the P_cLac143_ promoter system [20]) to regulate the expression of PmHAS or SeHasA, and integrated the two respective genes at the *acsA* locus of Syn7002 (Fig. 2A). Fully segregated strains (HA01 harboring *pmHAS* and HA03 containing *seHasA*), as well as C-terminal GFP-tagged versions (HA02 and HA04 respectively), were confirmed by colony PCR (Fig. 2B). Immunoblotting experiments confirmed that expression of both HA synthases was induced upon IPTG addition (Fig. 2C), with overexpression of PmHAS and SeHasA resulting in the appearance of either a 110 kDa or a 49 kDa protein, respectively. Overexpression of either HA synthase impacted cell growth, with SeHasA having a more severe effect (Fig. S1A).

**Figure 2.**
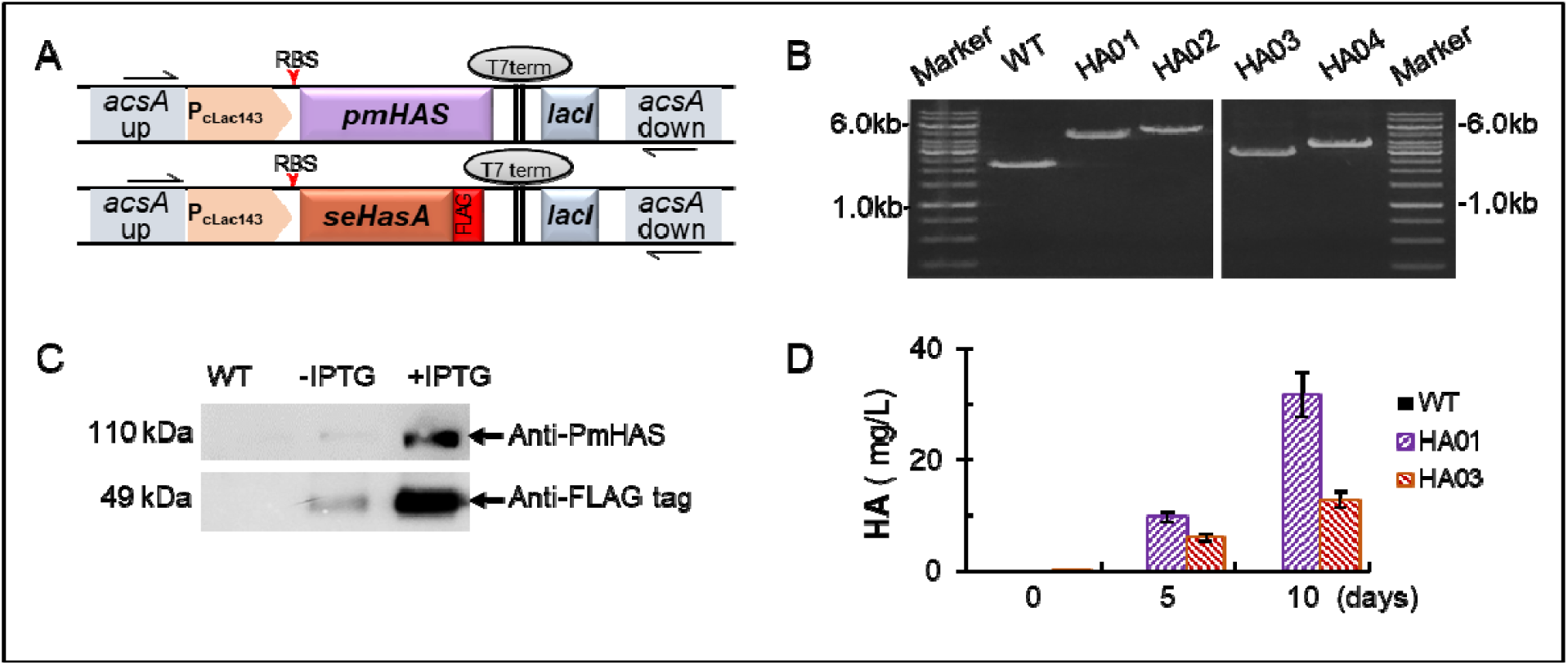
Introduction of two different HA synthases into Syn7002. **A.** Artificial operons used for expression of different HA synthases from the *acsA* locus of Syn7002, controlled by the inducible promoter P_cLac143_. **B.** Genomic DNA PCR analysis of strains with *pmHAS* (HA01) or *seHasA* (HA03), as well as their *sfGFP*-tagged version (HA02 and HA04), with specific primers listed in Table S1. Expected PCR sizes are 4773 bp for strain HA01, 5508 bp for strain HA02, 3279 bp for HA03, 3983 bp for HA04 and 2303 bp for the *acsA* WT locus. **C.** Western blot analysis of PmHAS or SeHasA expression in response to IPTG addition, using either anti-PmHAS or anti-FLAG antibodies, respectively. **D.** Quantification of HA in the growth medium in cultures induced with 1mM IPTG, at the time points indicated. Error bars represent standard deviation of three biological replicates (n=3), measured in duplicate. HA01: Δ*acsA*∷P_cLac143_-*pmHAS*; HA03: Δ*acsA*∷P_cLac143_-*seHasA*

HA synthase activity in these strains was evaluated by quantifying HA present in the medium after IPTG induction using a specific ELISA Kit, which uses a specific HA-binding protein for recognition and quantification of HA with molecular mass higher than 35 kDa [29]. While WT cultures produced no detectable HA, strains expressing either of the HA synthases produced moderate amounts of HA upon induction. For strain HA01, the HA concentration in the medium increased from 9.8±0.85 mg/L (at 5 days post-induction) to 31.9±4.0 mg/L (at 10 days post-induction) while strain HA03 had an overall lower HA production in comparison to the HA01 strain (Fig. 2B). However, as the higher HA concentration at 10 days was linked to culture decline, all subsequent HA production experiments were performed with cell cultures at an earlier growth phase (up to 5 days post-induction).

To understand better the reasons behind the low productivity of strain HA03 (expressing the integral membrane SeHasA synthase), localization of both enzymes was studied by adding a C-terminal superfolder-GFP (sfGFP) tag. This tag did not negatively affect HA synthesis activity, with similar amounts of HA detected in the growth medium 5 days post-IPTG induction (Fig. S1B). Confocal microscopy revealed that cells expressing sfGFP-tagged derivatives of PmHAS (HA02) and SeHasA (HA04) were larger than WT, with SeHasA expression having a more pronounced effect (Fig. S2). sfGFP fluorescence was concentrated in the cytoplasm of HA02 cells but mostly found in random patches in HA04. Thylakoid membranes in HA04 cells also seemed obviously distorted, possibly by mistargeting of integral membrane protein SeHasA or due to accumulation of HA in the lumen space. Consequently, all further engineered strains used PmHAS, as this soluble HA synthase was more benign and productive than its integral counterpart.

### 3.2. Characterization of HA produced by PmHAS in Syn7002

In addition to the specific ELISA assay, we employed several alternative methods to positively identify the polymers produced upon expressing PmHAS. Strain HA01 was used to isolate released cyanobacterial polysaccharides by ethanol precipitation from culture supernatants after 5 days of IPTG induction. The obtained material (HA01-HA) was analyzed by hyaluronidase digestion, HPLC-SEC, and LC-MS/MS. Hyaluronidase digestion of HA01-HA, similarly to that of commercial HA, resulted in a loss of detection by the ELISA assay and a shift to much smaller polymer sizes, as detected by HPLC-SEC (Fig. S3). LC-MS/MS analysis of the hyaluronidase-digested samples identified typical HA disaccharides with m/z 395 and tetrasaccharide with m/z 774.89 both in the digested HA01-HA sample and the commercial HA control (Fig. S4), further verifying the identity of the produced polymer.

### 3.3. Cellulose removal benefits HA production

The biosynthesis of HA in our modified Syn7002 strains relies solely on intermediates derived from photosynthetic carbon fixation (see Fig. 1), which are also commonly used by the cells for the generation of internal glycogen storage granules [30] or are converted to UDP-Glucose (UDP-Glc) for synthesis of cell wall components, such as cellulose or other types of polysaccharides [31]. The carbon flux towards HA synthesis is therefore potentially limited by these other pathways. As the biosynthesis pathways involved in cell wall polysaccharides have not been fully characterized in Syn7002, we focused on the possible effect of blocking cellulose synthesis or lowering the glycogen level on HA production. Therefore, several deletion mutants were constructed to investigate their influence on HA production (Fig. 3A).

**Figure 3.**
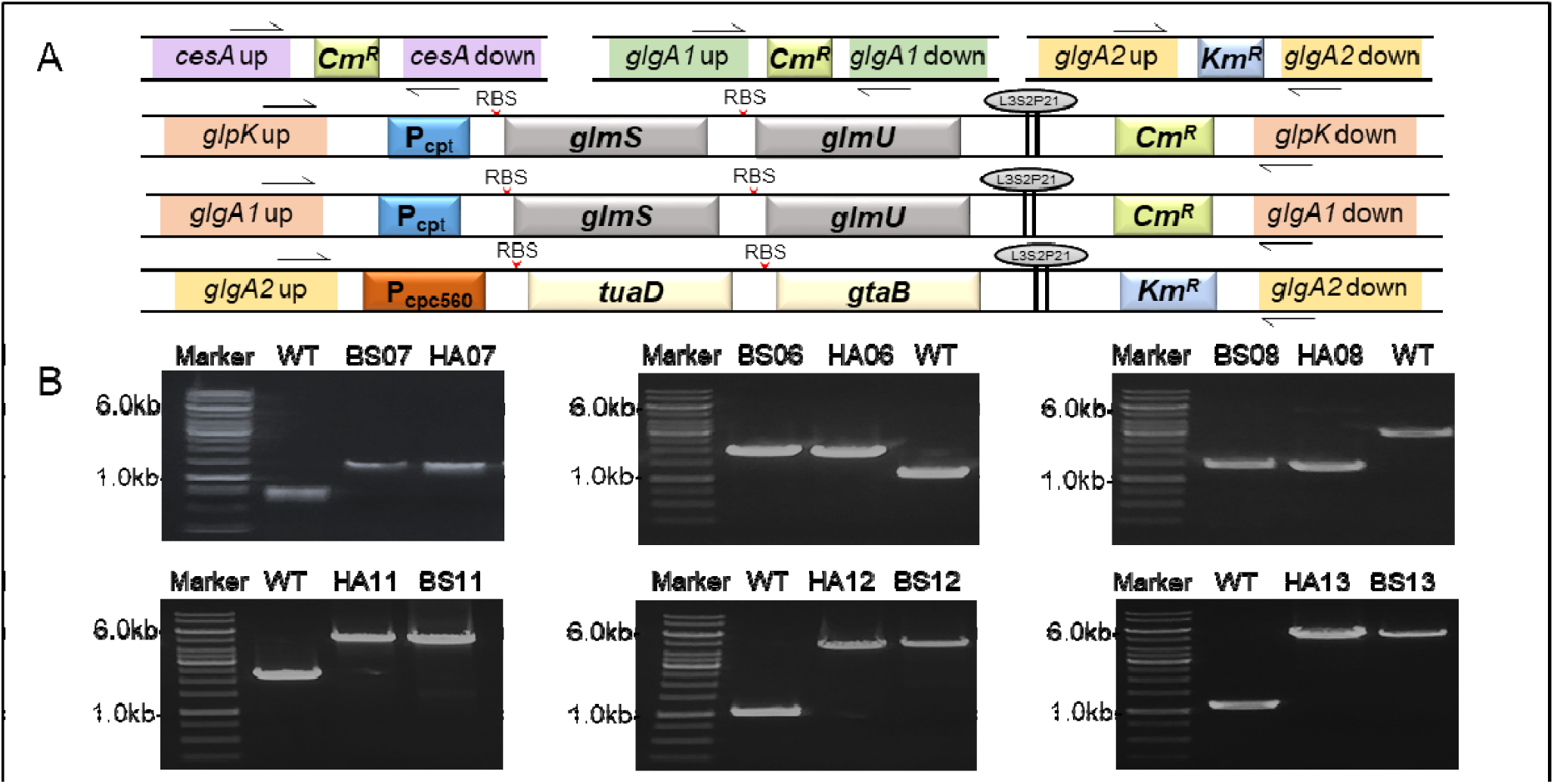
Overview of the different modifications for improved PmHAS-dependent HA production in Syn7002. **A.** Schematic images of constructs used for deletion of competing pathways and heterologous expression of precursor biosynthesis artificial operons. **B.** Gel images of gDNA PCR segregation tests for all the strains constructed in this work. BS06, BS07, BS08 are background strains, and HA06, HA07, HA08 are PmHAS-containing strains with *glgA2, glgA1* and *cesA* knock-out mutation. Expected PCR product sizes: 1259bp (BS07 and HA07), 1470bp (BS06 and HA06), 1365bp (BS08 and HA08). The corresponding expected PCR sizes in WT are 566bp (*glgA2*), 938bp (*glgA1*) and 2863bp (*cesA*). BS11, BS12, BS13 and their respective PmHAS-containing strains HA11, HA12, HA13 have heterologous precursor pathway operons inserted in different sites. Expected PCR fragment sizes are 4723bp (HA11), 4955bp (HA12) and 4199bp (HA13). Primers used for segregation tests are shown in Table S1.

Deletion of either of the two glycogen synthase genes (*glgA1* and *glgA2*) had only minor effects on HA production in comparison to the HA01 strain (Fig. S5), even though each deletion should increase glucose-1-phosphate pools and decrease glycogen biosynthesis by ≈50% [32]. The limited effects of partial glycogen depletion on HA production may imply that either Glc-1-P could not be efficiently converted to UDP-GlcUA or the limiting step for HA production might be related to the synthesis of the other precursor, UDP-GlcNAc. Cellulose, an insoluble polysaccharide that is also produced by some cyanobacteria, is synthesized in Syn7002 by a cellulose synthase (CesA), the deletion of which was shown to be essential for high-level cellulose production in Syn7002 [16]. Our results show that overexpression of PmHAS in strain HA08 (*cesA* knock-out background) led to a mild growth impairment in comparison to both WT and the basic HA01 strain, which was offset by substantially improved HA production, reaching 80.4±4.2 mg/L on the 5^th^ day post-induction (in comparison to 14.8±2.0mg/L in HA01, Fig. 4A). This marked improvement in HA production upon disruption of cellulose synthesis may be due to a higher flux from UDP-glucose to UDP-GlcUA, one of the HA precursors, as mentioned above. On the other hand, as cellulose was found to form a laminar layer in Syn7002, located in the vicinity of the peptidoglycan layer [16], there is also the possibility that endogenous cellulose removal may facilitate excretion of HA by removal of a physical barrier.

**Figure 4.**
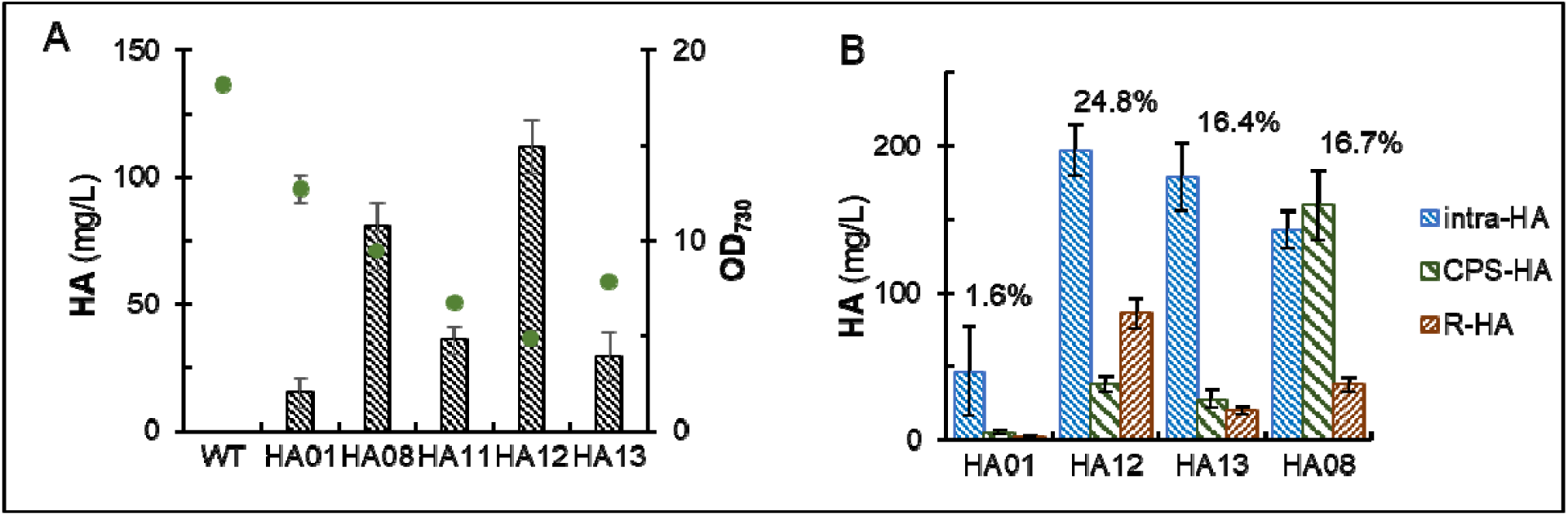
HA production by different strains. **A.** HA production (bar graphs) and cell density (dot plots) comparison at 5 days post IPTG induction. **B.** Assessment of HA excretion by HA quantification in different fractions. R-HA: HA released into growth medium; CPS-HA: capsular HA attached to cell surface; intra-HA: intracellular HA. Numbers in % represent the calculated photosynthetic carbon partitioning to total HA production. Error bars represent standard deviation of technical triplicates. HA01: Δ*acsA*∷P_cLac143_-*pmHAS*; HA08: Δ*cesA*+Δ*acsA*∷P_cLac143_-*pmHAS*; HA11: *glpK*∷P_cpt_*-glmS-glmU-Cm*^*R*^+Δ*acsA*∷P_cLac143_-*pmHAS*; HA12: Δ*glgA1*∷P_cpt_*-glmS-glmU-Cm*^*R*^+Δ*acsA*∷P_cLac143_-*pmHAS*; HA13: Δ*glgA2*∷P_cpc560_*-tuaD-gtaB-Km*^*R*^+Δ*acsA*∷P_cLac143_-*pmHAS*

### 3.4. HA yield is improved by overexpression of enzymes involved in precursor biosynthesis

A commonly utilized strategy in other bacterial strains engineered for HA production is to strengthen the biosynthesis of the precursors UDP-GlcUA and/or UDP-GlcNAc [33]. Studies using recombinant strains of *B. subtilis* overexpressing either SeHasA or PmHAS have shown that UDP-glucuronic acid (UDP-GlcUA) biosynthesis can be a limiting step for production of HA [14]. Conversely, the co-overexpression of PmHAS with GcaD (or GlmU), involved in synthesis of UDP-GlcNAc, produced very low molecular mass HA and improved HA titer to a lesser extent [12]. Although most of the enzymes involved in HA precursor biosynthesis have predicted homologs in cyanobacteria (as shown in Fig. 1), the expression, functionality and regulation of those Syn7002 gene products have so far not been studied and it was therefore preferable to manipulate precursor biosynthesis by expressing heterologous enzymes.

An artificial operon to improve UDP-GlcNAc biosynthesis, P_cpt_*-glmS-glmU* was introduced into either the *glpK* neutral locus in strain HA11 [18] or the *glgA1* site in strain HA12 (Fig. 3). HA production increased in both strains, to 111.9±10.6 mg/L in HA12 and 35.9±5.5 mg/L in HA11 (Fig. 4A), demonstrating that enhanced biosynthesis of UDP-GlcNAc improves HA yields in Syn7002, an effect magnified when glycogen synthesis was simultaneously inhibited. Although HA production was not improved by merely deleting *glgA1* (or *glgA2*), the combination of lower glycogen biosynthesis with increasing synthesis of the precursor UDP-GlcNAc did successfully divert carbon flow to the final product, HA. Separately, to increase UDP-GlcUA precursor pools, the P_cpc560_-*tuaD*-*gtaB* operon was introduced into the other glycogen synthase locus (*glgA2*) of Syn7002 in strain HA13 (Fig. 3). HA production (29.3±9.5 mg/L) improved in comparison to HA01, albeit to a lesser extent than in strain HA12 (Fig. 4A). Interestingly, overexpression of UDP-GlcUA-related enzymes in Syn7002 did not influence HA titers as strongly as in heterotrophic organisms [14]. As heterologous HA production diverts intermediates away from cell wall biogenesis, growth was affected by HA production, with higher HA titers generally resulting in lesser growth (lower OD_730_), particularly in the case of strain HA12, which had both the lowest cell density and the highest titer (Fig. 4A).

### 3.5. Released HA is only a minor fraction of the total production

As the mechanism for HA secretion in PmHAS-utilizing bacterial strains is unknown, we tested the excretion efficiency of producing strains by quantifying HA isolated from different locations. We found that in all the strains tested large amounts of HA were retained within the cells, ranging from 42% to roughly 88%. Strain HA12, with the UDP-GlcNAc pathway enhanced, was able to release a higher portion of HA into the medium, however the vast majority of the product (about 60%) was still trapped within the cell (Fig. 4B). Interestingly, strain HA08 had the highest amount of capsular HA among all the strains. Nearly 50% of the total HA production for strain HA08 was found attached to the cell surface (Fig. 4B). One possible explanation is that removal of the cellulose layer facilitates secretion of HA, even if it is not completely released from the cell surface. In light of these results, it seems clear that, even though all the generated strains improved their HA production capabilities, secretion and release of such large polymer is still a major bottleneck. Clearly, the mechanism responsible for polysaccharide (HA) excretion in cyanobacteria is unable to efficiently export the large amount of HA produced by the engineered strains, which may also be one of the causes for the growth defects observed.

Total HA production of these strains were calculated by summing up the values obtained from different fractions and photosynthetic CO_2_ partitioning was estimated according to measured dry cell biomass, carbon fraction in HA of 44.4% and total cellular carbon as 51.3% of cell biomass (as previously described [34]).Strain HA12 has the highest carbon partitioning efficiency (≈25%), a substantial increase in comparison to strain HA01 (converting <2% of fixed carbon to HA). Overexpression of GlmS and GlmU most likely diverts carbon flow from the primary photosynthetic intermediate fructose-6-phosphate, to synthesize UDP-GlcNAc, resulting in comparatively better carbon partitioning (Fig. 4B and Table S3), though at the expense of cell growth (Fig. 4A). Strain HA08 (with *cesA* deleted) showed a similar CO_2_ partitioning to HA as strain HA13 (roughly 16%), but with a better growth performance, which may be more suitable for sustainable production of HA.

### 3.6. The molecular weight of excreted HA varies with different pathway modifications

Size-exclusion chromatography was used to characterize the molecular mass of the excreted polymers produced by the different strains, by comparison to 2-2.2 MDa commercial HA and 1 MDa PEG standard as our reference polymers. All the samples from different strains had two major peaks, one with a molecular mass greater than 2-2.2 MDa and a smaller one with mass greater than 1 MDa, referred as “UHMW-HA” and “HMW-HA” respectively (Fig. 5). The peaks detected were eluted and further analyzed by HA quantification (Fig. 5B) and FTIR (Fig. S6). The quantification of the HA content of each eluted peak generally fits with the peak intensity shown by the chromatography spectra, indicating that the introduced modifications, while possibly influencing precursor levels, resulted in changes to the size of produced HA. As shown in Fig. 5, strains HA08 and HA12 had a higher proportion of the UHMW-HA, whereas strains HA01 and HA13 strains excreted mainly HMW-HA. The reason for this is at the moment unclear but might be related to the different regulation mechanisms at the level of polymerization and secretion. FTIR spectra for the major chromatography peaks isolated from the HA-producing strains, but not from WT Syn7002, were very similar to that of commercial HA (Fig. S6), again confirming the identity of the produced polymers.

**Figure 5.**
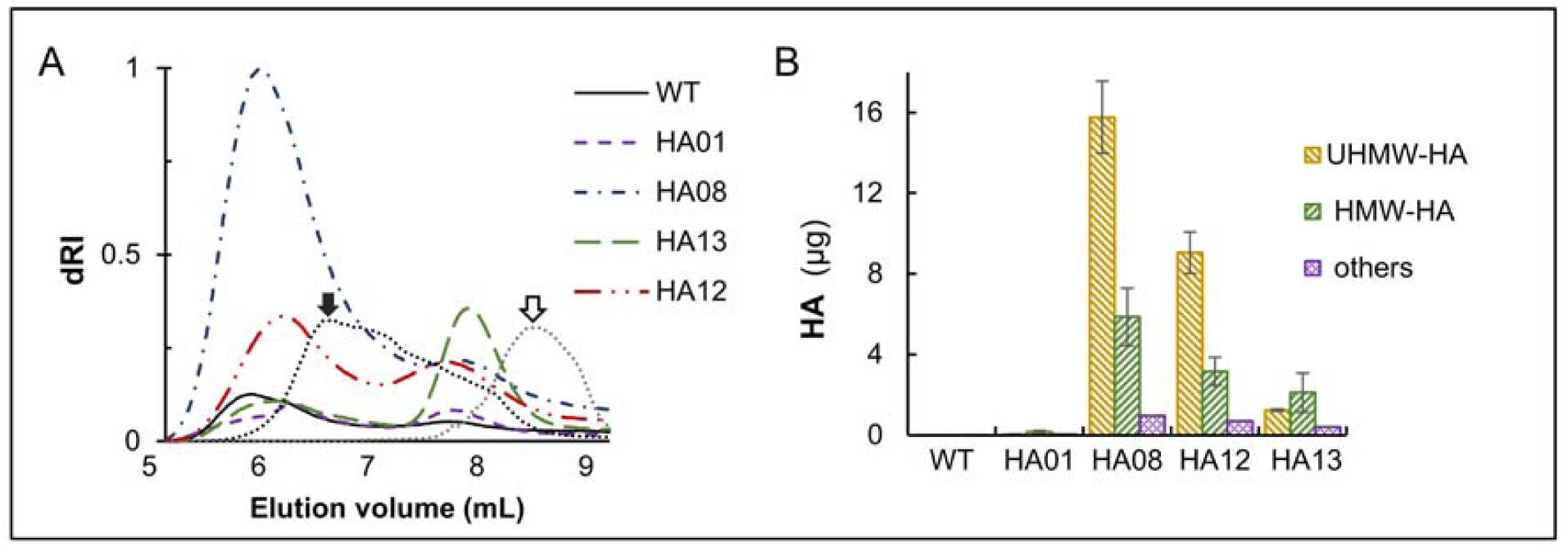
Size estimation of HA polymers from different strain by HPLC-SEC. **A.** Chromatograms of released polymers from different HA producing strains. Black filled arrow: commercial HA (molecular mass 2-2.2MDa); black outlined arrow: PEG polymer standard (molecular mass 1 MDa). **B.** HA content in each eluted peak. Peaks were eluted according to the intensity changes of dRI detector (elution times may vary between samples of different HA producing strains). Error bars represent standard deviation of technical duplicates.

## 4. Conclusions

This work shows that cyanobacteria are promising hosts for the production of biomedically-relevant sugar polymers solely by photosynthesis, paving the way for sustainable production of hyaluronic acid. Heparosan was recently shown to be produced by heterologous expression of a heparosan synthase in *Synechococcus* sp. PCC 7942, although with an extremely low yield (3 µg/L) [5]. The best performing strain in the current report (HA12) shows a several thousand-fold improvement in comparison to the heparosan-producing strains, perhaps owing to codon-optimization of the introduced enzymes, more robust growth of our cyanobacterial host in comparison to *Synechococcus* sp. PCC7942 and iterative metabolic engineering. HA production was successfully improved by using different strategies, such as deleting the endogenous cellulose synthase gene and overexpressing precursor biosynthesis enzymes and, especially in the case of the UDP-GlcNAc pathway, combination of precursor biosynthesis with glycogen depletion was particularly beneficial for production, though at the cost of growth performance.

Even if these results are encouraging, several issues still remain to be solved before cyanobacteria can compete with the yields observed in heterotrophic hosts. Increasing the photosynthetic performance of cyanobacteria is a commonly used strategy for improved production yield, as this could improve up-stream precursor pools for HA synthesis and other metabolites [35]. PmHAS-mediated HA secretion may be further improved by manipulation of other competing pathways (such as O-antigen or peptidoglycan synthesis), which could also lead to a further improvement of HA excretion, by removal of physical impairments.

As environmentally-conscious processes become increasingly relevant in relation to the observed climate changes, development of efficient engineered cyanobacterial cell factories will allow biopolymer production bypassing competition for food-grade materials, arable land or freshwater resources, thus contributing to a more sustainable bio-economy.

## Acknowledgements

This work was supported by NTU grants M4080306 to BN and M4081714 to PJN. The authors would like to thank Dr. Wahyu Surya (SBS, NTU) for assistance with size-exclusion chromatography analysis of samples and Associate Professor Jaume Torres (SBS, NTU) for usage of the FTIR spectrometer. We would also like to thank the NTU Optical Microscopy for cell imaging analysis and NTU Phenomics Centre for LC-MS/MS analysis of HA fragments. The authors are grateful to Prof. Bertil Andersson for his constant support and encouragement and his pivotal role in establishing the CyanoSynBio@NTU laboratory.

## Contributions

LZ, TTS, PJN and BN conceptualized and designed the present study. TTS performed the initial cloning of HA synthases in the expression vector, LZ performed all other strain engineering, cyanobacteria cultivation, production tests and product analysis. All authors contributed to data analysis, manuscript drafting and revision, agree to authorship and approve the final manuscript for submission.

## Supplementary materials

**Table S1.** List of primers used in this work.

**Table S2.** List of plasmids constructed in this work.

**Table S3.** Estimation of total fixed carbon partitioning to HA in different producing strains.

**Figure S1.** Introduction and expression of two different HA synthases into Syn7002.

**Figure S2.** Confocal microscopy images of Syn7002 cells expressing HA synthase.

**Figure S3.** Hyaluronidase digestion of cyanobacterial HA compared to commercial HA.

**Figure S4.** LC-MS/MS analysis of hyaluronidase digested samples from strain HA01 and commercial HA.

**Figure S5.** FTIR spectra of commercial HA and HPLC-purified HA produced by different strains.

## Abbreviations

WT: wild type
GFP: Green Fluorescent Protein
RBS: ribosome binding site
UDP: uridine diphosphate
IPTG: Isopropyl β-D-1-thiogalactopyranoside
PEG: polyethylene glycol

