## Supplementary figures and images for "Photosynthetic conversion of CO_2_ to hyaluronic acid by engineered cyanobacteria"

### Figures S1-S6 and Tables S1-S3

#### Slide 1
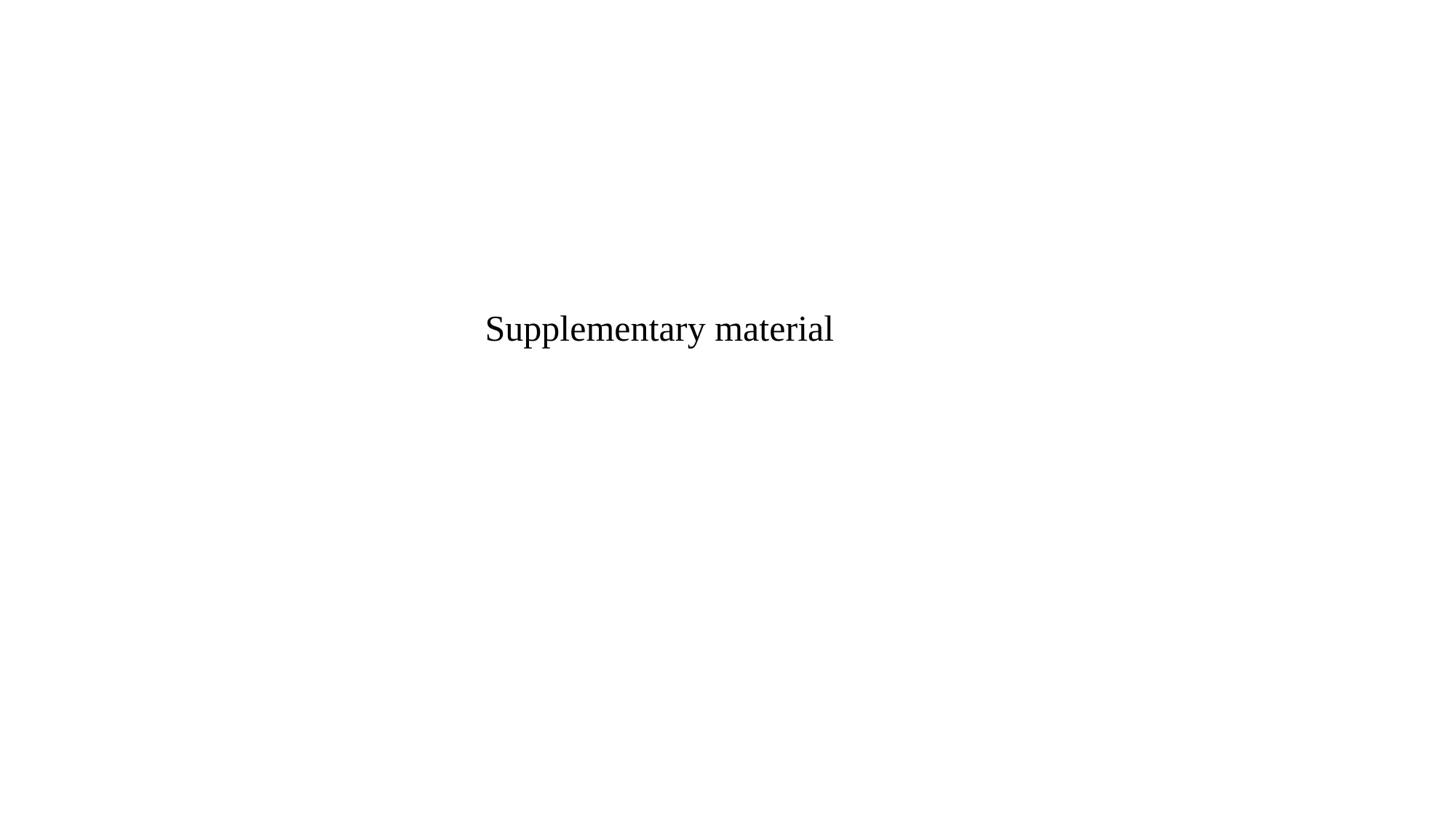

#### Slide 2
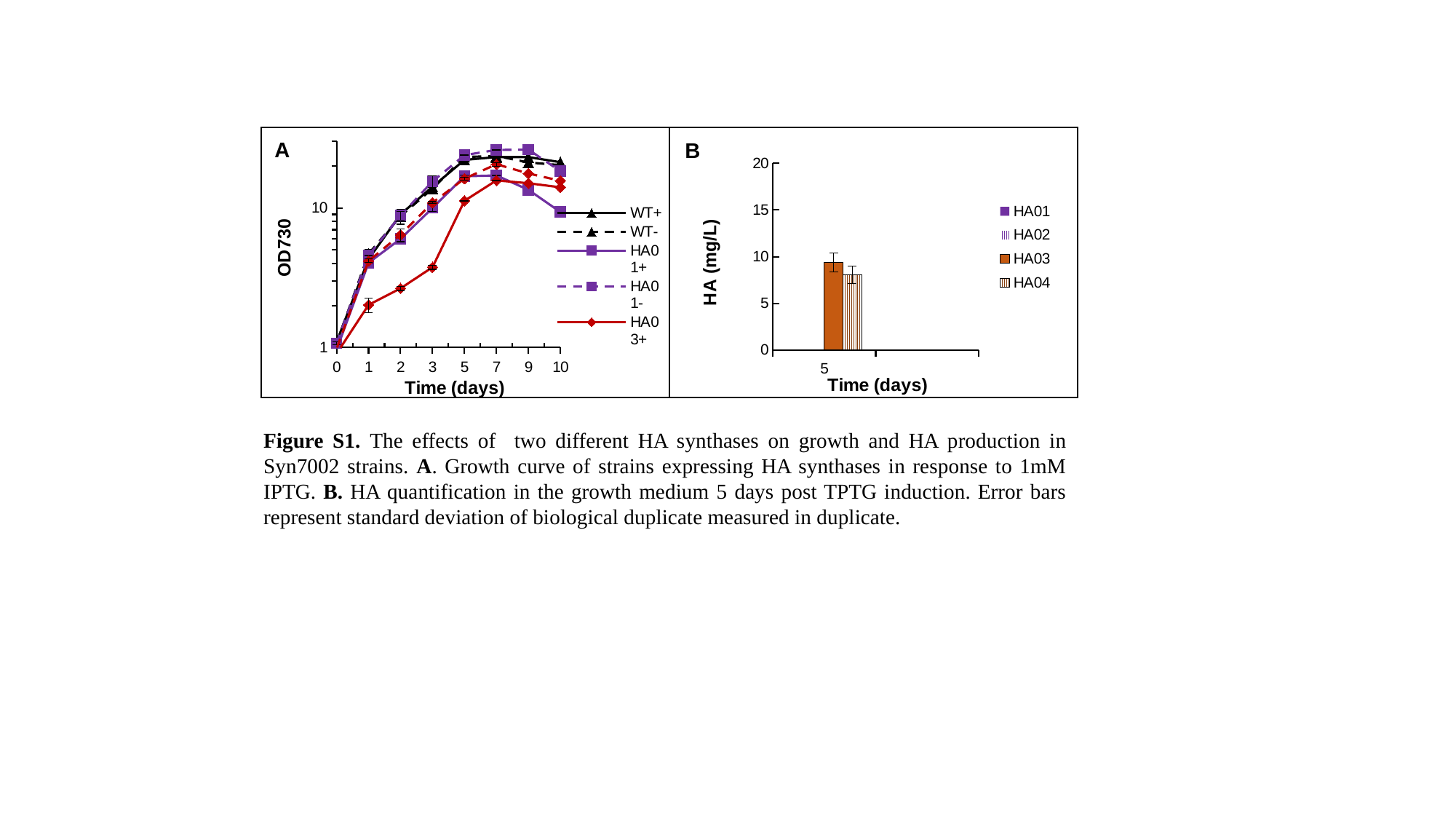

#### Slide 3
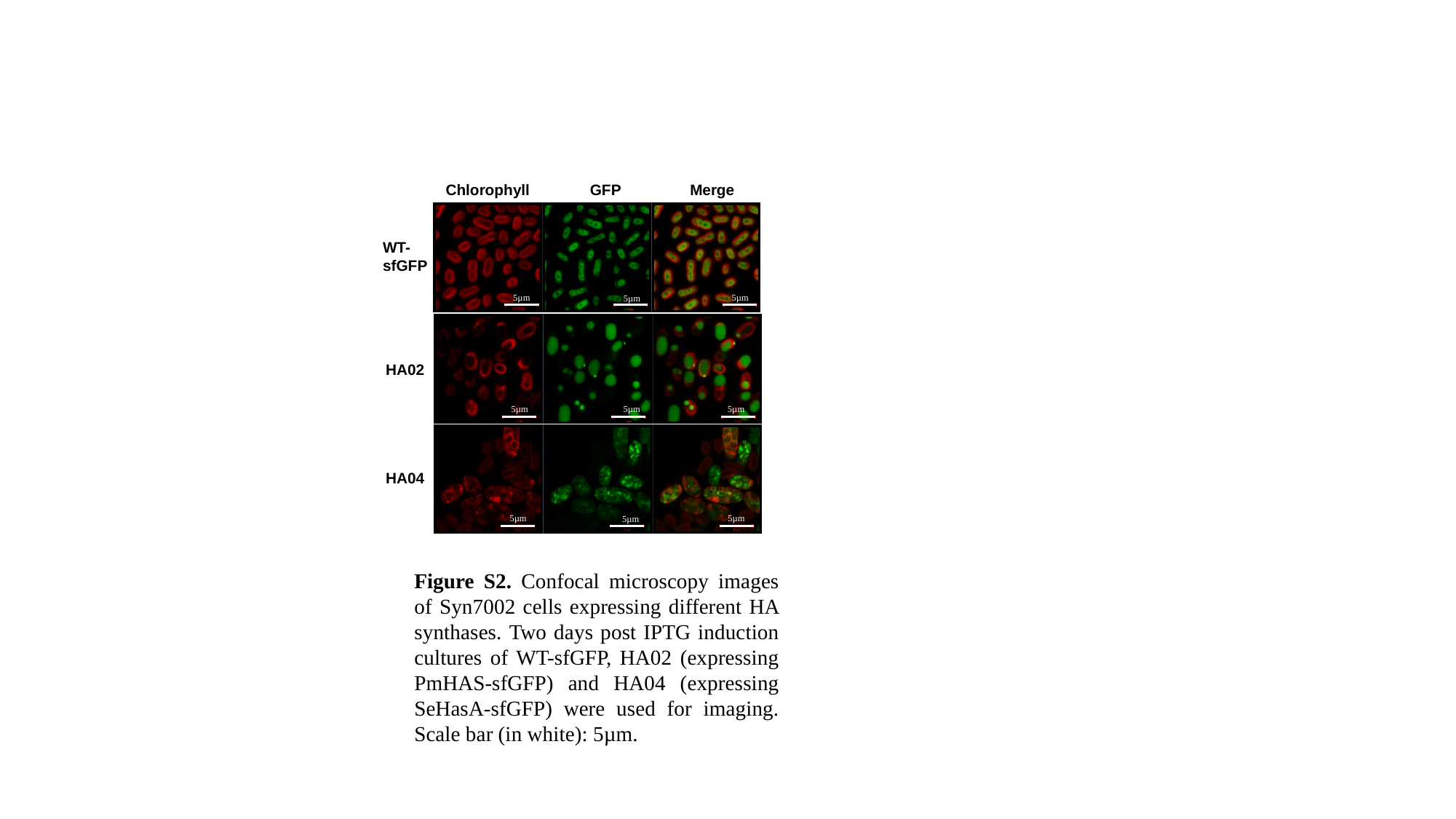

#### Slide 4
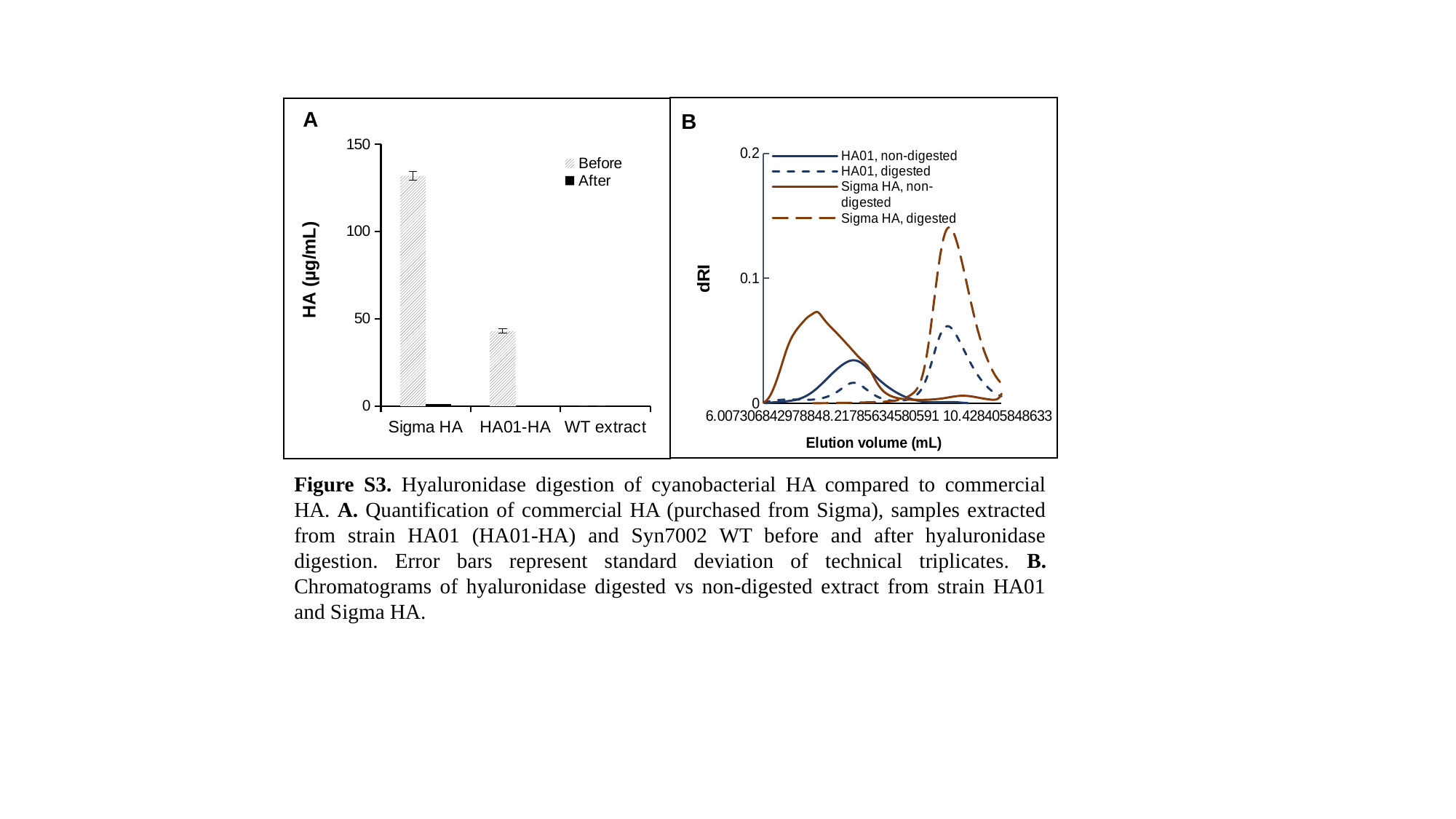

#### Slide 5
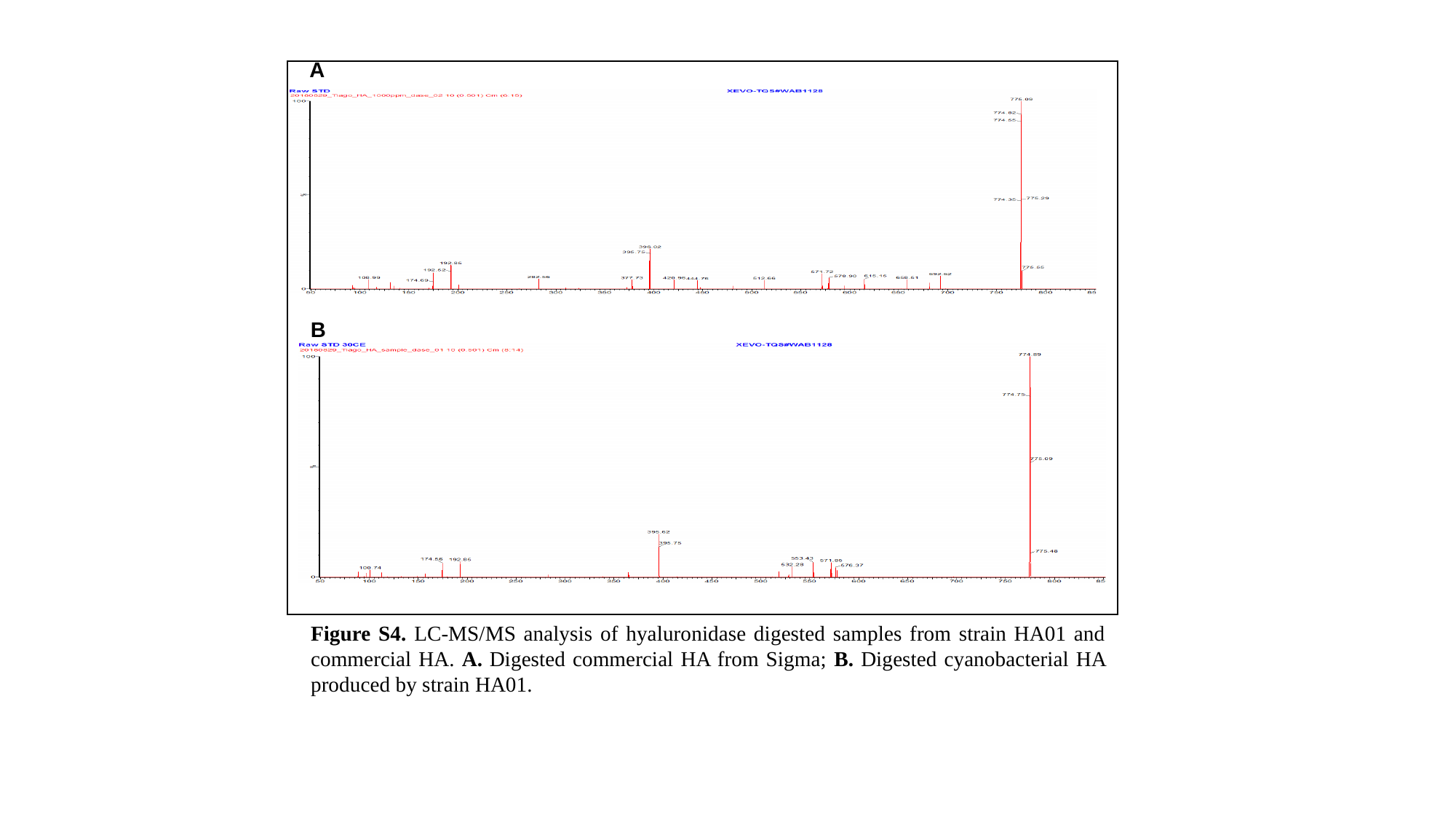

#### Slide 6
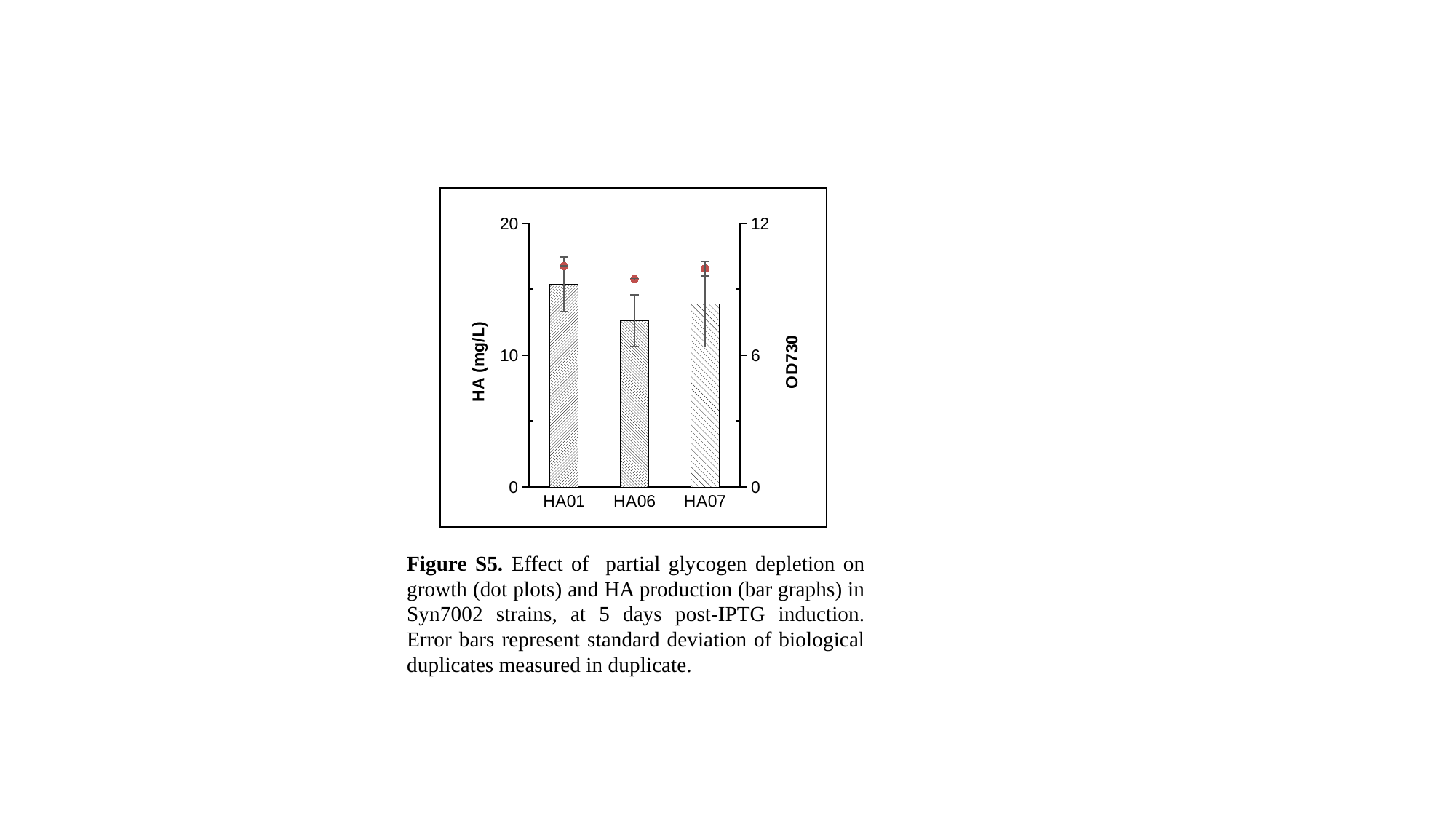

#### Slide 7
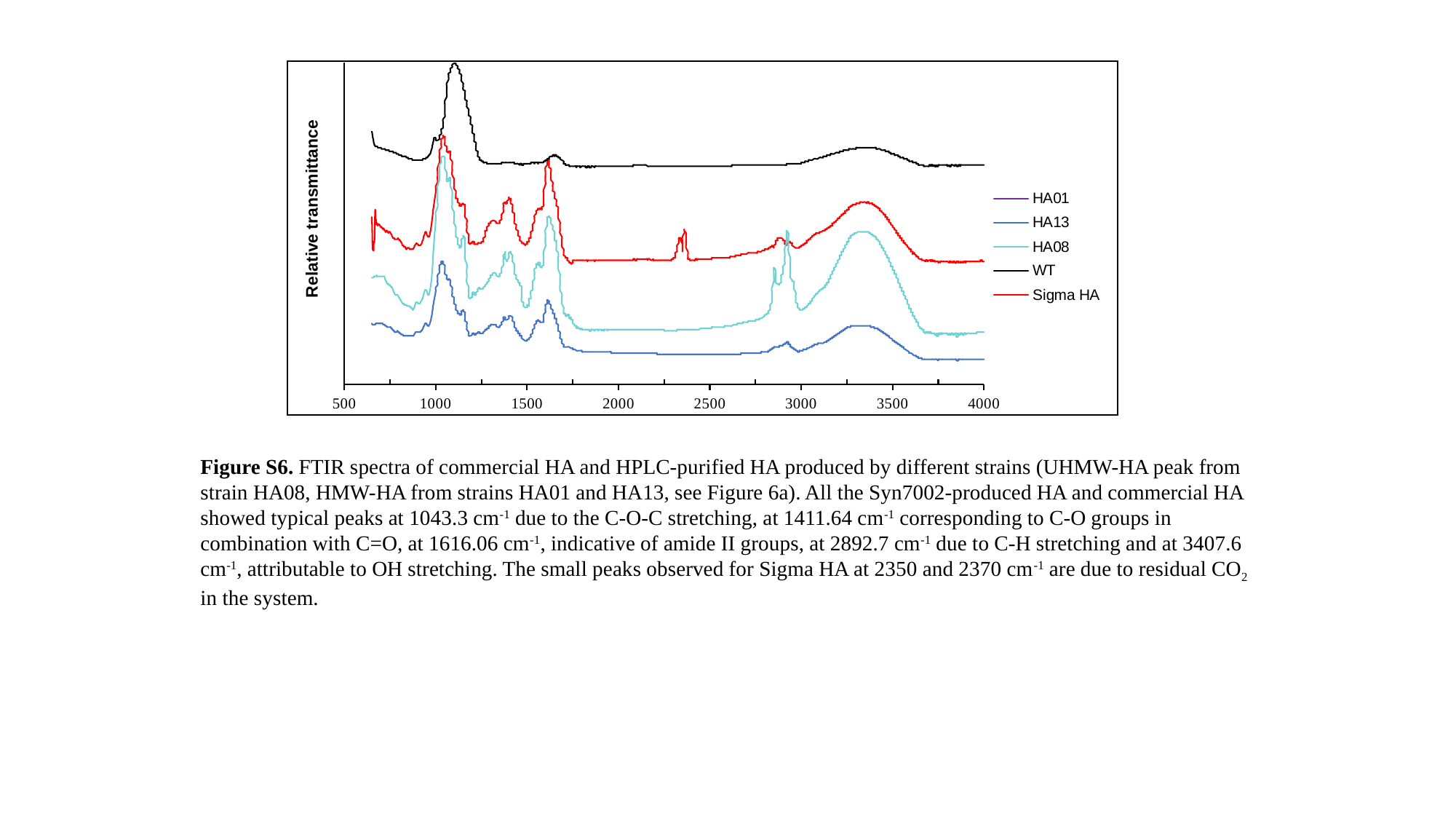

#### Slide 8
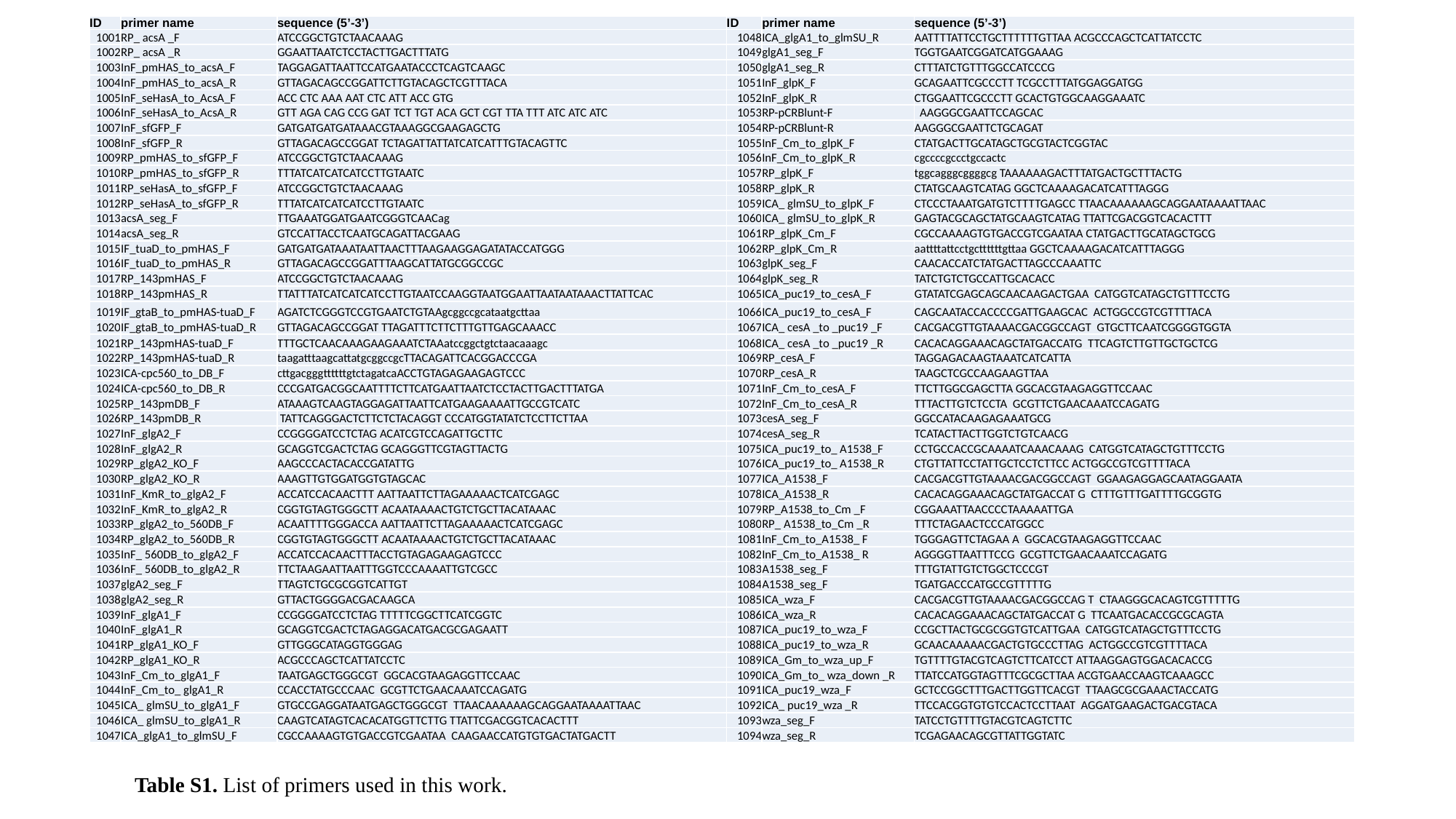

#### Slide 9
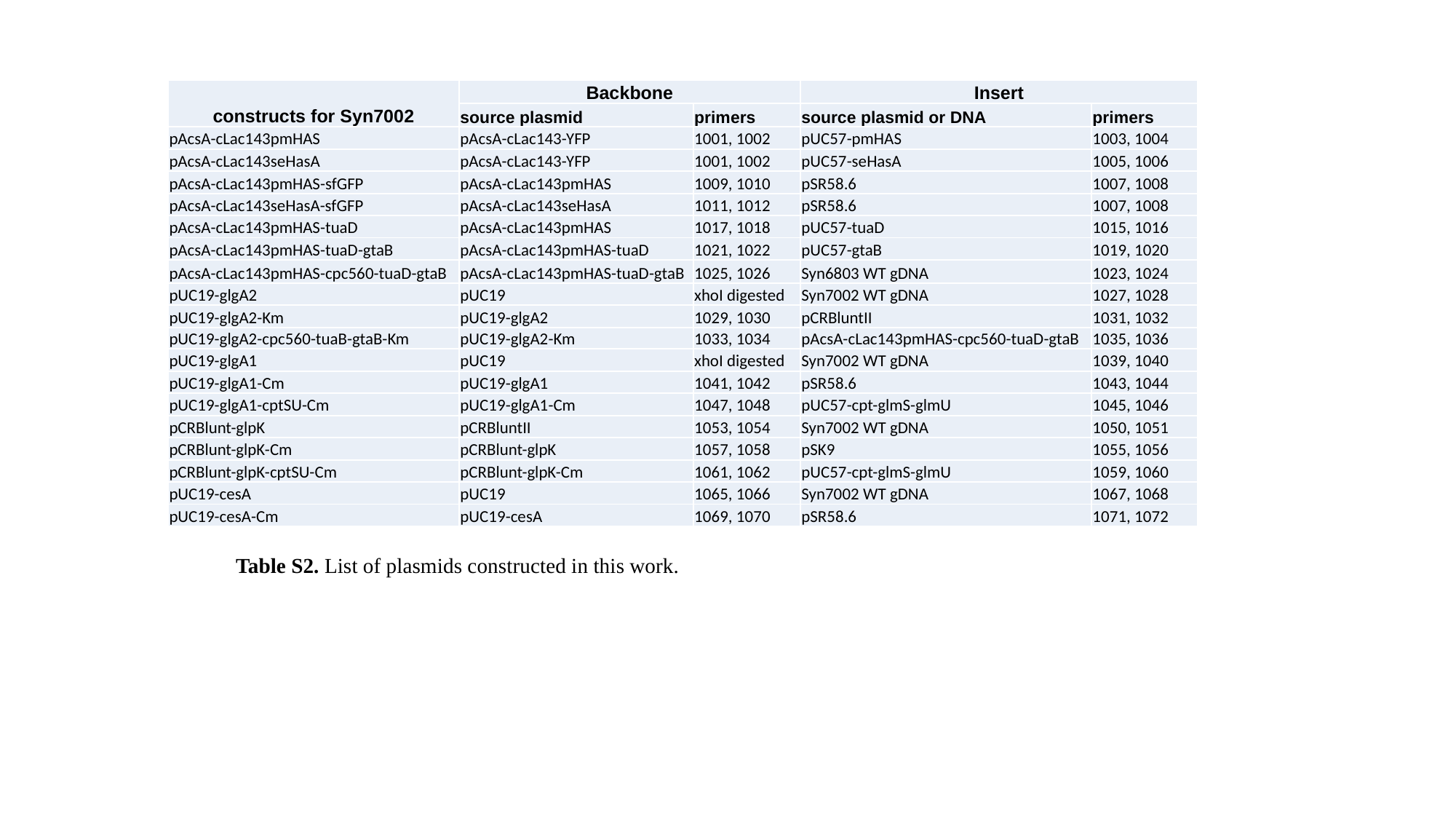

#### Slide 10
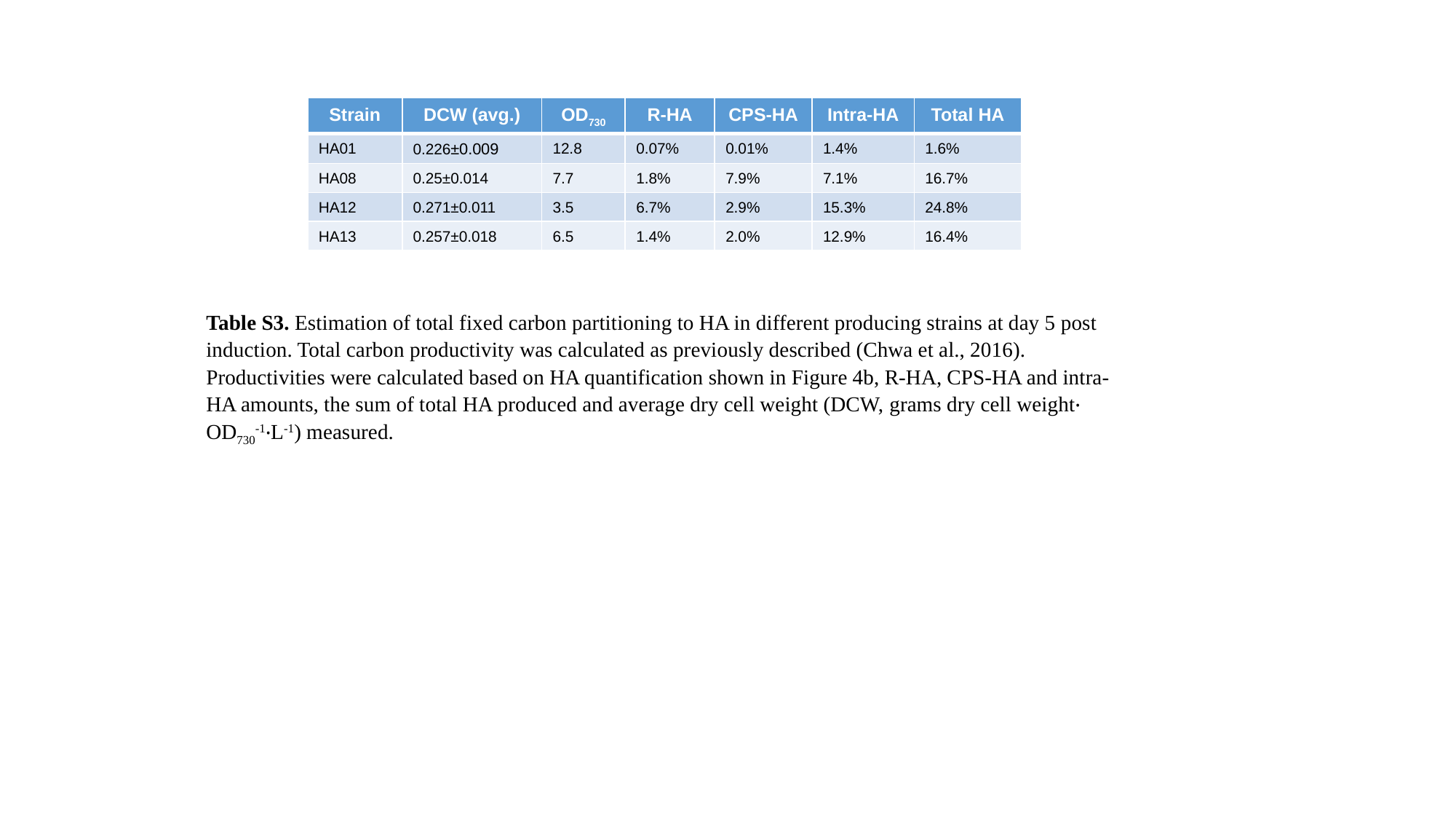
